# Insects and associated arthropods analyzed during medicolegal death investigations in Harris County, Texas, USA: January 2013-April 2016

**DOI:** 10.1101/071027

**Authors:** Michelle R. Sanford

## Abstract

The application of insect and arthropod information to medicolegal death investigations is one of the more exacting applications of entomology. Historically limited to homicide investigations, the integration of full time forensic entomology services to the medical examiner’s office in Harris County has opened up the opportunity to apply entomology to a wide variety of manner of death classifications and types of scenes to make observations on a number of different geographical and species-level trends in Harris County, Texas, USA. In this study, a retrospective analysis was made of 203 forensic entomology cases analyzed during the course of medicolegal death investigations performed by the Harris County Institute of Forensic Sciences in Houston, TX, USA from January 2013 through April 2016. These cases included all manner of death classifications, stages of decomposition and a variety of different scene types that were classified into decedents transported from the hospital (typically associated with myiasis or sting allergy; 3.0%), outdoor scenes (32.0%) or indoor scenes (65.0%). Ambient scene air temperature at the time scene investigation was the only significantly different factor observed between indoor and outdoor scenes with average indoor scene temperature being slightly cooler (25.2°C) than that observed outdoors (28.0°C). Relative humidity was not found to be significantly different between scene types. Most of the indoor scenes were classified as natural (43.3%) whereas most of the outdoor scenes were classified as homicides (12.3%). All other manner of death classifications came from both indoor and outdoor scenes. Several species were found to be significantly associated with indoor scenes as indicated by a binomial test, including *Blaesoxipha plinthopyga* (Sarcophagidae), all Sarcophagidae including *B*. *plinthopyga*, *Megaselia scalaris* (Phoridae), *Synthesiomyia nudiseta* (Muscidae) and *Lucilia cuprina* (Calliphoridae). The only species that was a significant indicator of an outdoor scene was *Lucilia eximia* (Calliphoridae). All other insect species that were collected in five or more cases were collected from both indoor and outdoor scenes. A species list with month of collection and basic scene characteristics with the length of the estimated time of colonization is also presented. The data presented here provide valuable casework related species data for Harris County, TX and nearby areas on the Gulf Coast that can be used to compare to other climate regions with other species assemblages and to assist in identifying new species introductions to the area. This study also highlights the importance of potential sources of uncertainty in preparation and interpretation of forensic entomology reports from different scene types.

## INTRODUCTION

While forensic entomology has a broad scope, one of its most challenging applications is to medicolegal death investigations. The application of the information provided by insects and associated arthropods to death investigations can take many forms ranging from direct association with cause of death (e.g. insect sting allergy), to information related to decedent travel history (e.g. ticks traveling with their host decedents), to the estimation of time of insect colonization (TOC) that can be used to approximate the post-mortem interval (PMI; [1]). The use of insect colonization and development in estimating the PMI is one of the most well-known applications of forensic entomology in the medicolegal setting [2].

Since its first recorded application to the death investigation of the infamous bloody sickle [3] to the more recent application to the time of death of Danielle van Dam [4] forensic entomology has had a strong association with homicide investigation and crime scene analysis [1,2,5]. The recent addition of forensic entomology services to the medical examiner’s office has led to an even broader application of forensic entomology to medicolegal death investigations across all manners of death [6]. When one starts to investigate the insects associated with a broad range of decedents, scenes and manners of death one starts to appreciate the breadth of forensic entomology casework opportunities. One of the first steps to understanding these cases is in the identification of the insects involved in colonizing human remains in a given geographic location, their seasonality and the characteristics of the scenes where they are found.

Forensic entomology surveys of different geographic locations are often performed as a first step in establishing baseline data (e.g. [7–9]). However, these types of studies are often based on trapping adult flies or using animal models and are not typically based on casework information. In Texas, survey information specific to forensically important insects is rare with only a few published surveys available from Central Texas [9,10]. Blow fly (Diptera: Calliphoridae) trapping studies were prevalent prior to and immediately following the screwworm eradication program but the focus of these efforts was mainly to detect *Cochliomyia hominivorax* [11–13]. A series of experiments examining the interaction between the newly introduced *Chrysomya* sp. into the United States also recorded the presence of several blow flies in Central parts of Texas with traps as well [14,15] but the focus of these studies was not primarily for survey purposes.

Species records and trends related to casework have been published in other geographic areas. In North America these types of surveys have included the Hawaiian island of Oahu [16,17] and Western Canada [18]. These types of data are rarely published but are exceedingly important to understanding the local fauna important to casework in different geographic locations especially when practitioners may be reliant upon old and restricted taxonomic keys, which may not include all the species local to the area. To date no published data relating casework and the local forensically important insects in Texas or specifically in Harris County has been recorded. In this study the species, trends and seasonality of the forensically important insects associated with casework are reported for Harris County, Texas, USA. These data will aid in not only guiding identification and tracking of new species in the area but also in guiding research questions into the species of importance in this area. This study also allows for comparison with previous studies, particularly with respect to larger scale trends in insect colonization patterns such as indoor and outdoor scenes, seasonality and introduced species.

## MATERIALS AND METHODS

### Casework

Cases involving insect or related arthropod specimens collected and analyzed during the course of medicolegal death investigations performed by the Harris County Institute of Forensic Sciences (HCIFS) as dictated under Texas Code of Criminal Procedure 49.25 [19], for the period of January 2013 through April 2016 (N = 203) were included in this dataset. These cases involved the collection of specimens at the scene or during the autopsy and occasionally at both locations (Figure 1). The procedure for specimen collection follows standard operating procedures developed for the office. Briefly, this includes collection of representative insect and/or related arthropod specimens from representative locations on the body and scene representing the oldest identified life stages associated with the body. Specimens are divided into representative portions, preserved by the most appropriate method (e.g. hot water kill followed by 70% ethanol for larval fly specimens [20]) and reared to the adult stage, if possible, for confirmation of identification.

**Figure 1:**
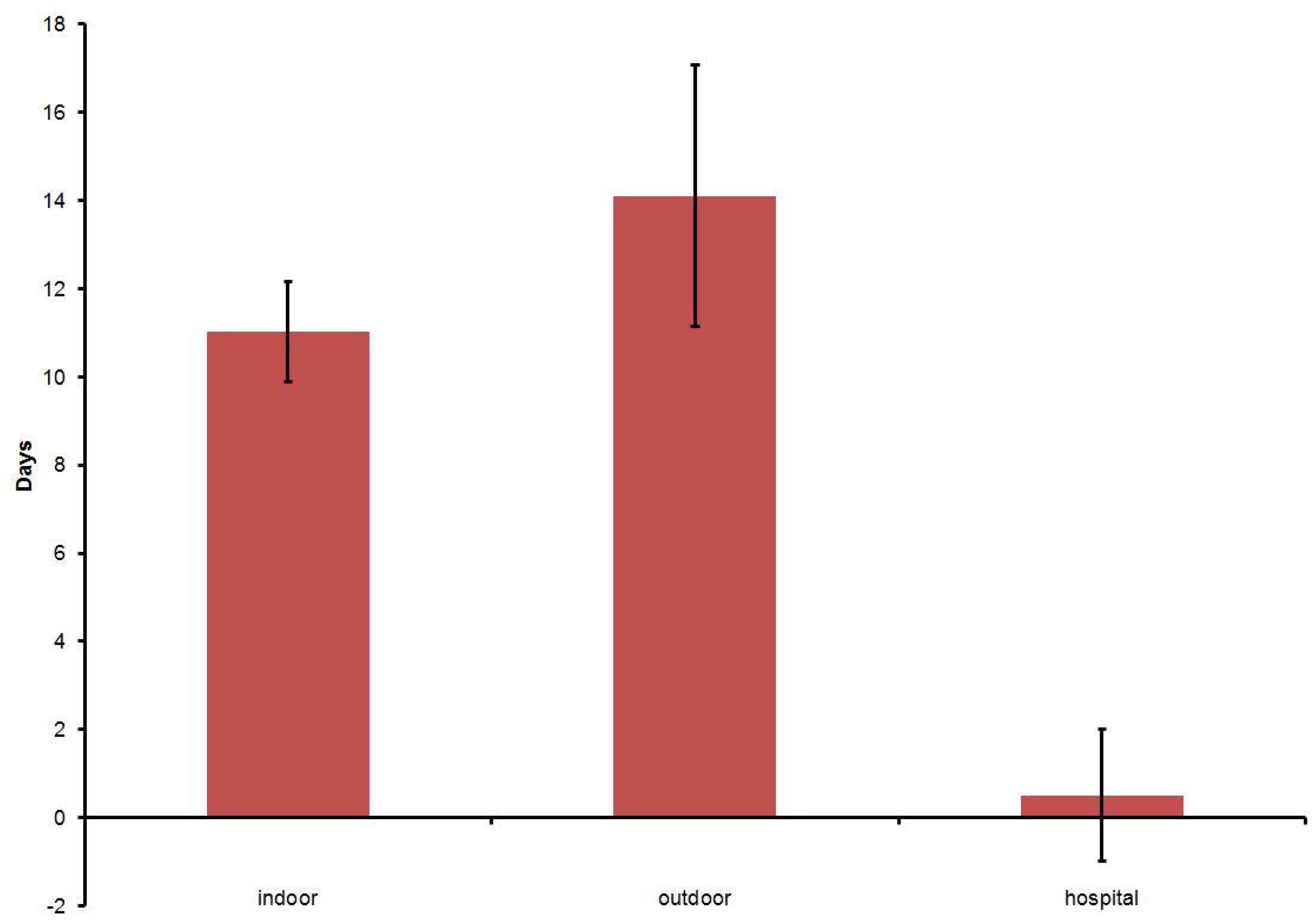
Forensic Entomology Cases by Official Manner of Death and Scene Location. Percentage of forensic entomology cases over the study period (January 2013 - April 2016) by official manner of death classification, showing the proportion of cases where the scene was located indoors (blue) or outdoors (red). (N=203).

Identification of specimens is based on morphology using life stage and taxon appropriate taxonomic keys and literature, which are referenced in each individual report and the most frequently used publications are presented in Supplemental Table 1. Specialized keys and consultation with taxonomic experts are consulted as needed depending upon the specific case. Dr. T. Whitworth (Washington State University) also generously provided the HCIFS with a small adult reference blow fly (Diptera: Calliphoridae) collection in 2013 to facilitate identification of adult Calliphoridae specimens.

Cases that require only identification of specimens, such as for stinging insect cases, may not require further analysis, but a majority of cases require further analysis to estimate the age of the collected specimens and to provide a time of colonization (TOC) estimate. Larval Diptera and Coleoptera specimens are measured for length and each collected species is photographed for representative identifying features using a Keyence VHX-600 digital microscope with an attached VHX-S15E used as needed to stack images to enhance depth of field (Keyence Corp., Itasca, IL, USA). The estimated age of the larvae is calculated using the accumulated degree hours (ADH) method [1,21] and taxon appropriate published development data for each species using either published length data and/or development stage. Many scenes were located on private property with access granted by law enforcement. No protected species were collected. The completed TOC estimate and report for the specimens are included with the full autopsy report for each case involving the collection of entomology specimens.

During the course of scene investigation, temperature and humidity recordings are made using a combined thermometer-hygrometer (Fisher Scientific Education™, S66279, Fisher Scientific, Pittsburgh, PA, USA). Scene conditions such as location of the body with respect to shade or sun outdoors or thermostat settings indoors are also recorded to assist in the assessment of temperature modifications at the scene that might influence differences in temperature from the nearest weather station. Data is collected for ADH calculations from the nearest National Oceanic and Atmospheric Administration (NOAA) quality controlled weather station for each scene.

### Statistics

Basic statistics were calculated for the forensic entomology cases where insects were collected from 2013 through April of 2016 (two cases were excluded as they reported on the absence of insects in the only two burial cases over the 3+ year period) for a total of 203 cases. Basic statistics (e.g. mean, standard deviation, count) were calculated for scene location (indoor, outdoor or hospital), month of scene investigation, length of TOC estimate, ambient scene temperature (℃), ambient scene relative humidity and the presence of insect species collected for each case (Table 1). Statistical comparisons of cases and case related factors (e.g. temperature differences by case type and species presence indoors vs. outdoors) were calculated using a Kruskal-Wallis Χ^2^ test for continuous data (e.g. ambient scene temperature, relative humidity and TOC estimate length) or an exact binomial test for categorical data (e.g. scene location) for species which were collected in five or more cases. All statistical calculations and tests were computed using Microsoft Excel^®^ and R 3.2.4 (R Team 2016) using the R Commander package [23].

**Table 1.**
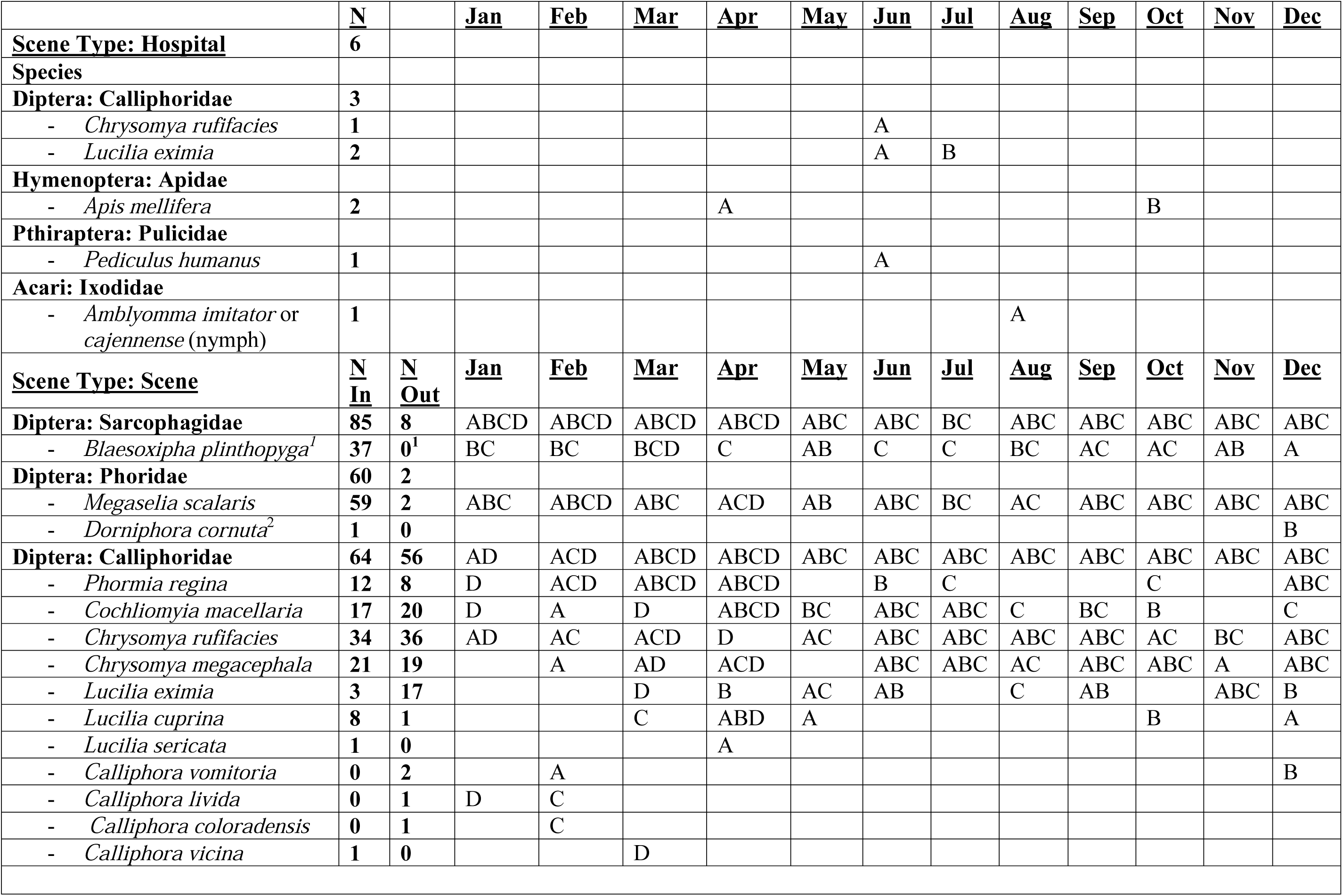
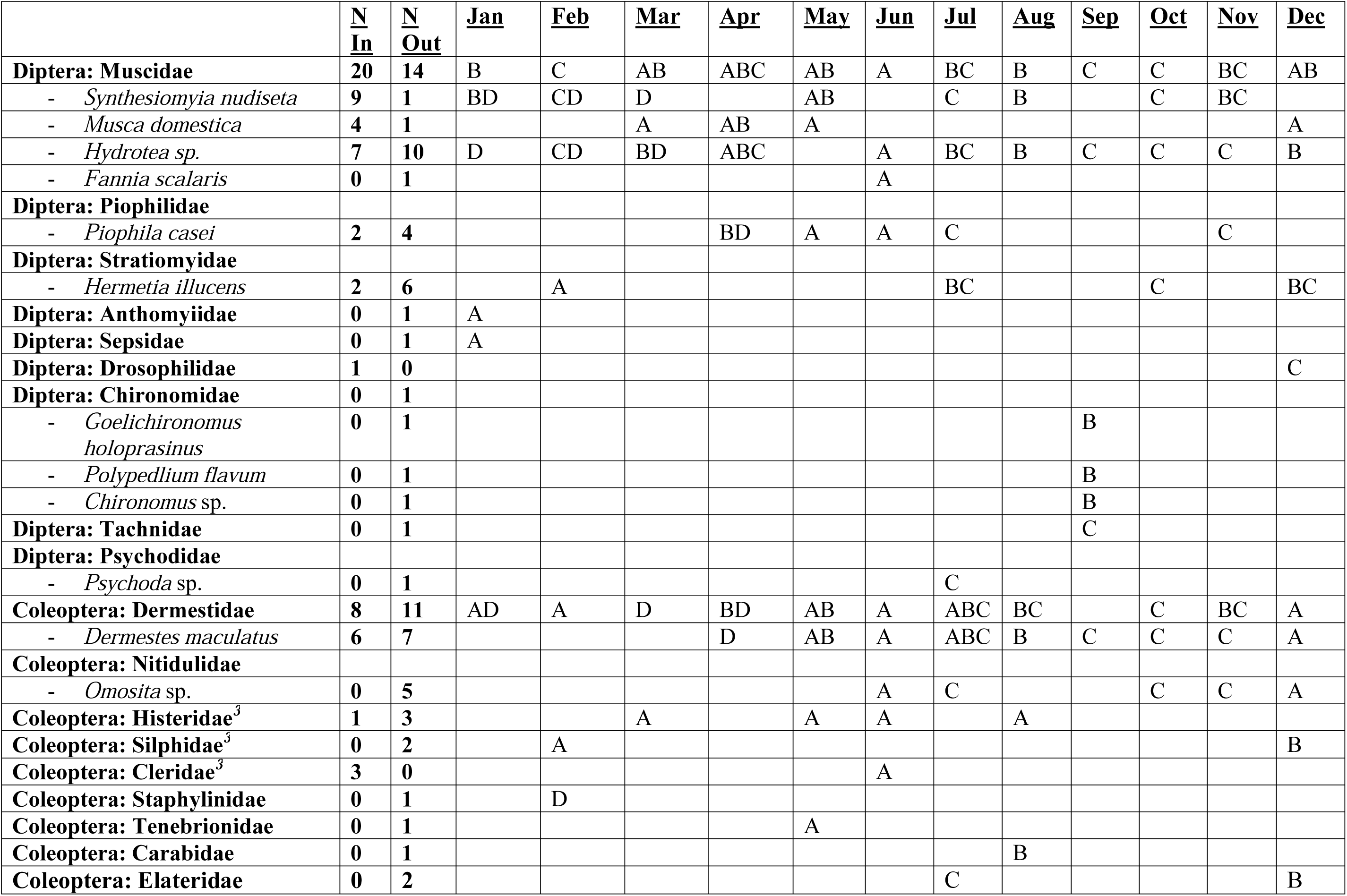
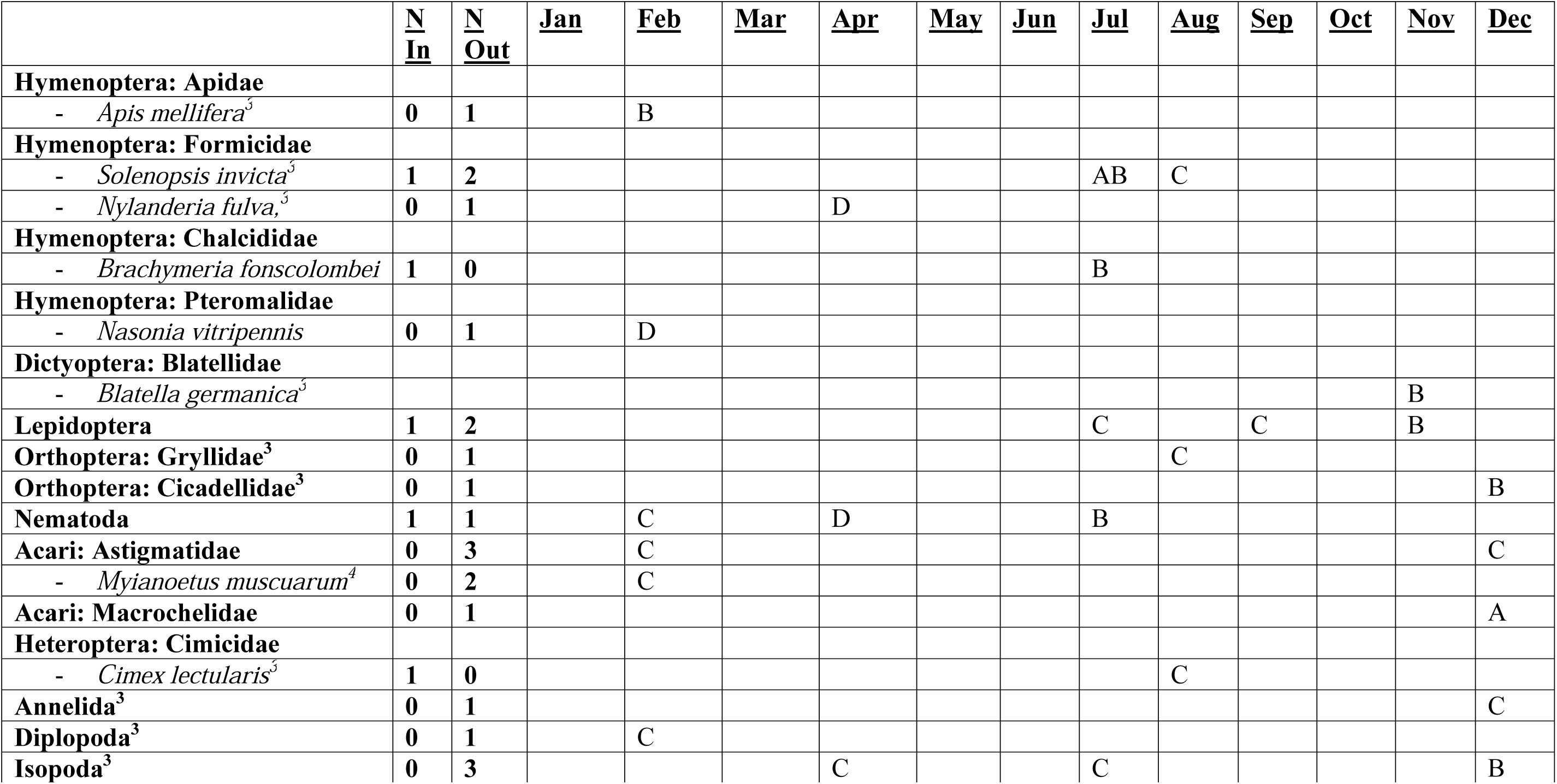
A list of insect and arthropod species collected during selected medicolegal death investigations handled by the Harris County Institute of Forensic Sciences from January 2013 through April 2016. The month of recorded collection is indicated by the letters A for 2013, B for 2014, C for 2015 and D for 2016. ^1^Confirmation of this species is based on the appearance of male genitalia from reared adult males and this could be an underestimate of actual abundance for this species because reared males may not always be obtained. ^2^Identification of this specimen is based on a single larva and is tentatively identified to the species level. ^3^These species may be regularly observed at scenes but not regularly collected and reported upon because they are not typically related to time of colonization estimation. ^4^This species was recently associated with *Synthesiomyia nudiseta* [39].

|  | <u>N</u> |  | <u>Jan</u> | <u>Feb</u> | <u>Mar</u> | <u>Apr</u> | <u>May</u> | <u>Jun</u> | <u>Jul</u> | <u>Aug</u> | <u>Sep</u> | <u>Oct</u> | <u>Nov</u> | <u>Dec</u> |
| --- | --- | --- | --- | --- | --- | --- | --- | --- | --- | --- | --- | --- | --- | --- |
| **Scene Type: Hospital** | **6** |  |  |  |  |  |  |  |  |  |  |  |  |  |
| **Species** |  |  |  |  |  |  |  |  |  |  |  |  |  |  |
| **Diptera: Calliphoridae** | **3** |  |  |  |  |  |  |  |  |  |  |  |  |  |
| - *Chrysomya rufifacies* | **1** |  |  |  |  |  |  | A |  |  |  |  |  |  |
| - *Lucilia eximia* | **2** |  |  |  |  |  |  | A | B |  |  |  |  |  |
| **Hymenoptera: Apidae** |  |  |  |  |  |  |  |  |  |  |  |  |  |  |
| - *Apis mellifera* | **2** |  |  |  |  | A |  |  |  |  |  | B |  |  |
| **Pthiraptera: Pulicidae** |  |  |  |  |  |  |  |  |  |  |  |  |  |  |
| - *Pediculus humanus* | **1** |  |  |  |  |  |  | A |  |  |  |  |  |  |
| **Acari: Ixodidae** |  |  |  |  |  |  |  |  |  |  |  |  |  |  |
| - *Amblyomma imitator* or *cajennense* (nymph) | **1** |  |  |  |  |  |  |  |  | A |  |  |  |  |
| **Scene Type: Scene** | <u>N</u> | <u>N</u> | <u>Jan</u> | <u>Feb</u> | <u>Mar</u> | <u>Apr</u> | <u>May</u> | <u>Jun</u> | <u>Jul</u> | <u>Aug</u> | <u>Sep</u> | <u>Oct</u> | <u>Nov</u> | <u>Dec</u> |
|  | <u>In</u> | <u>Out</u> |  |  |  |  |  |  |  |  |  |  |  |  |
| **Diptera: Sarcophagidae** | **85** | **8** | ABCD | ABCD | ABCD | ABCD | ABC | ABC | BC | ABC | ABC | ABC | ABC | ABC |
| - *Blaesoxipha plinthopyga*<sup>1</sup> | **37** | **0**<sup>1</sup> | BC | BC | BCD | C | AB | C | C | BC | AC | AC | AB | A |
| **Diptera: Phoridae** | **60** | **2** |  |  |  |  |  |  |  |  |  |  |  |  |
| - *Megaselia scalaris* | **59** | **2** | ABC | ABCD | ABC | ACD | AB | ABC | BC | AC | ABC | ABC | ABC | ABC |
| - *Dorniphora cornuta*<sup>2</sup> | **1** | **0** |  |  |  |  |  |  |  |  |  |  |  | B |
| **Diptera: Calliphoridae** | **64** | **56** | AD | ACD | ABCD | ABCD | ABC | ABC | ABC | ABC | ABC | ABC | ABC | ABC |
| - *Phormia regina* | **12** | **8** | D | ACD | ABCD | ABCD |  | B | C |  |  | C |  | ABC |
| - *Cochliomyia macellaria* | **17** | **20** | D | A | D | ABCD | BC | ABC | ABC | C | BC | B |  | C |
| - *Chrysomya rufifacies* | **34** | **36** | AD | AC | ACD | D | AC | ABC | ABC | ABC | ABC | AC | BC | ABC |
| - *Chrysomya megacephala* | **21** | **19** |  | A | AD | ACD |  | ABC | ABC | AC | ABC | ABC | A | ABC |
| - *Lucilia eximia* | **3** | **17** |  |  | D | B | AC | AB |  | C | AB |  | ABC | B |
| - *Lucilia cuprina* | **8** | **1** |  |  | C | ABD | A |  |  |  |  | B |  | A |
| - *Lucilia sericata* | **1** | **0** |  |  |  | A |  |  |  |  |  |  |  |  |
| - *Calliphora vomitoria* | **0** | **2** |  | A |  |  |  |  |  |  |  |  |  | B |
| - *Calliphora livida* | **0** | **1** | D | C |  |  |  |  |  |  |  |  |  |  |
| - *Calliphora coloradensis* | **0** | **1** |  | C |  |  |  |  |  |  |  |  |  |  |
| - *Calliphora vicina* | **1** | **0** |  |  | D |  |  |  |  |  |  |  |  |  |

Table 1. A list of insect and arthropod species collected during selected medicolegal death investigations handled by the Harris County Institute of Forensic Sciences from January 2013 through April 2016.
|  | <u>N</u><br><u>In</u> | <u>N</u><br><u>Out</u> | <u>Jan</u> | <u>Feb</u> | <u>Mar</u> | <u>Apr</u> | <u>May</u> | <u>Jun</u> | <u>Jul</u> | <u>Aug</u> | <u>Sep</u> | <u>Oct</u> | <u>Nov</u> | <u>Dec</u> |
| --- | --- | --- | --- | --- | --- | --- | --- | --- | --- | --- | --- | --- | --- | --- |
| **Diptera: Muscidae** | **20** | **14** | B | C | AB | ABC | AB | A | BC | B | C | C | BC | AB |
| - *Synthesiomyia nudiseta* | **9** | **1** | BD | CD | D |  | AB |  | C | B |  | C | BC |  |
| - *Musca domestica* | **4** | **1** |  |  | A | AB | A |  |  |  |  |  |  | A |
| - *Hydrotea* sp. | **7** | **10** | D | CD | BD | ABC |  | A | BC | B | C | C | C | B |
| - *Fannia scalaris* | **0** | **1** |  |  |  |  |  | A |  |  |  |  |  |  |
| **Diptera: Piophilidae** |  |  |  |  |  |  |  |  |  |  |  |  |  |  |
| - *Piophila casei* | **2** | **4** |  |  |  | BD | A | A | C |  |  |  | C |  |
| **Diptera: Stratiomyidae** |  |  |  |  |  |  |  |  |  |  |  |  |  |  |
| - *Hermetia illucens* | **2** | **6** |  | A |  |  |  |  | BC |  |  | C |  | BC |
| **Diptera: Anthomyiidae** | **0** | **1** | A |  |  |  |  |  |  |  |  |  |  |  |
| **Diptera: Sepsidae** | **0** | **1** | A |  |  |  |  |  |  |  |  |  |  |  |
| **Diptera: Drosophilidae** | **1** | **0** |  |  |  |  |  |  |  |  |  |  |  | C |
| **Diptera: Chironomidae** | **0** | **1** |  |  |  |  |  |  |  |  |  |  |  |  |
| - *Goelichironomus holoprasinus* | **0** | **1** |  |  |  |  |  |  |  |  | B |  |  |  |
| - *Polypedilium flavum* | **0** | **1** |  |  |  |  |  |  |  |  | B |  |  |  |
| - *Chironomus* sp. | **0** | **1** |  |  |  |  |  |  |  |  | B |  |  |  |
| **Diptera: Tachnidae** | **0** | **1** |  |  |  |  |  |  |  |  | C |  |  |  |
| **Diptera: Psychodidae** |  |  |  |  |  |  |  |  |  |  |  |  |  |  |
| - *Psychoda* sp. | **0** | **1** |  |  |  |  |  |  | C |  |  |  |  |  |
| **Coleoptera: Dermestidae** | **8** | **11** | AD | A | D | BD | AB | A | ABC | BC |  | C | BC | A |
| - *Dermestes maculatus* | **6** | **7** |  |  |  | D | AB | A | ABC | B | C | C | C | A |
| **Coleoptera: Nitidulidae** |  |  |  |  |  |  |  |  |  |  |  |  |  |  |
| - *Omosita* sp. | **0** | **5** |  |  |  |  |  | A | C |  |  | C | C | A |
| **Coleoptera: Histeridae**<sup>3</sup> | **1** | **3** |  |  | A |  | A | A |  | A |  |  |  |  |
| **Coleoptera: Silphidae**<sup>3</sup> | **0** | **2** |  | A |  |  |  |  |  |  |  |  |  | B |
| **Coleoptera: Cleridae**<sup>3</sup> | **3** | **0** |  |  |  |  |  | A |  |  |  |  |  |  |
| **Coleoptera: Staphylinidae** | **0** | **1** |  | D |  |  |  |  |  |  |  |  |  |  |
| **Coleoptera: Tenebrionidae** | **0** | **1** |  |  |  |  | A |  |  |  |  |  |  |  |
| **Coleoptera: Carabidae** | **0** | **1** |  |  |  |  |  |  |  | B |  |  |  |  |
| **Coleoptera: Elateridae** | **0** | **2** |  |  |  |  |  |  | C |  |  |  |  | B |
| **Hymenoptera: Apidae** |  |  |  |  |  |  |  |  |  |  |  |  |  |  |
| - *Apis mellifera*<sup>3</sup> | 0 | 1 |  | B |  |  |  |  |  |  |  |  |  |  |
| **Hymenoptera: Formicidae** |  |  |  |  |  |  |  |  |  |  |  |  |  |  |
| - *Solenopsis invicta*<sup>3</sup> | 1 | 2 |  |  |  |  |  |  | AB | C |  |  |  |  |
| - *Nylanderia fulva*,<sup>3</sup> | 0 | 1 |  |  |  | D |  |  |  |  |  |  |  |  |
| **Hymenoptera: Chalcididae** |  |  |  |  |  |  |  |  |  |  |  |  |  |  |
| - *Brachymeria fonscolombei* | 1 | 0 |  |  |  |  |  |  | B |  |  |  |  |  |
| **Hymenoptera: Pteromalidae** |  |  |  |  |  |  |  |  |  |  |  |  |  |  |
| - *Nasonia vitripennis* | 0 | 1 |  | D |  |  |  |  |  |  |  |  |  |  |
| **Dictyoptera: Blatellidae** |  |  |  |  |  |  |  |  |  |  |  |  |  |  |
| - *Blatella germanica*<sup>3</sup> |  |  |  |  |  |  |  |  |  |  |  |  | B |  |
| **Lepidoptera** | 1 | 2 |  |  |  |  |  |  | C |  | C |  | B |  |
| **Orthoptera: Gryllidae**<sup>3</sup> | 0 | 1 |  |  |  |  |  |  |  | C |  |  |  |  |
| **Orthoptera: Cicadellidae**<sup>3</sup> | 0 | 1 |  |  |  |  |  |  |  |  |  |  |  | B |
| **Nematoda** | 1 | 1 |  | C |  | D |  |  | B |  |  |  |  |  |
| **Acari: Astigmatidae** | 0 | 3 |  | C |  |  |  |  |  |  |  |  |  | C |
| - *Myianoetus muscuarum*<sup>4</sup> | 0 | 2 |  | C |  |  |  |  |  |  |  |  |  |  |
| **Acari: Macrochelidae** | 0 | 1 |  |  |  |  |  |  |  |  |  |  |  | A |
| **Heteroptera: Cimicidae** |  |  |  |  |  |  |  |  |  |  |  |  |  |  |
| - *Cimex lectularis*<sup>3</sup> | 1 | 0 |  |  |  |  |  |  |  | C |  |  |  |  |
| **Annelida**<sup>3</sup> | 0 | 1 |  |  |  |  |  |  |  |  |  |  |  | C |
| **Diplopoda**<sup>3</sup> | 0 | 1 |  | C |  |  |  |  |  |  |  |  |  |  |
| **Isopoda**<sup>3</sup> | 0 | 3 |  |  |  | C |  |  | C |  |  |  |  | B |

## RESULTS AND DISCUSSION

### Casework

Perhaps the most important thing to consider when examining these data is that they represent casework and not the results of planned ecological study, which imposes certain limitations on the statistical analyses completed and the conclusions drawn. Of the 203 cases analyzed, six were from hospitals without an associated scene investigation (3.0%), 65 were from outdoor scenes (32.0%) and 132 were from indoor scenes (65.0%). Scene type appears to play a significant role in not only the species of insects encountered (Table 1) but in the conditions one may encounter during scene investigation (Table 2; Figure 1). It is also important to note that the scene conditions reported here do not necessarily represent the conditions experienced by the colonizing insects because these data were collected when insects were collected after some period of development. These data represent the conditions in which these insects might be found during the course of routine death investigations, which may be valuable to understanding the conditions under which insects colonize human remains; a goal of applied forensic entomology as it is used in casework.

**Table 2.**
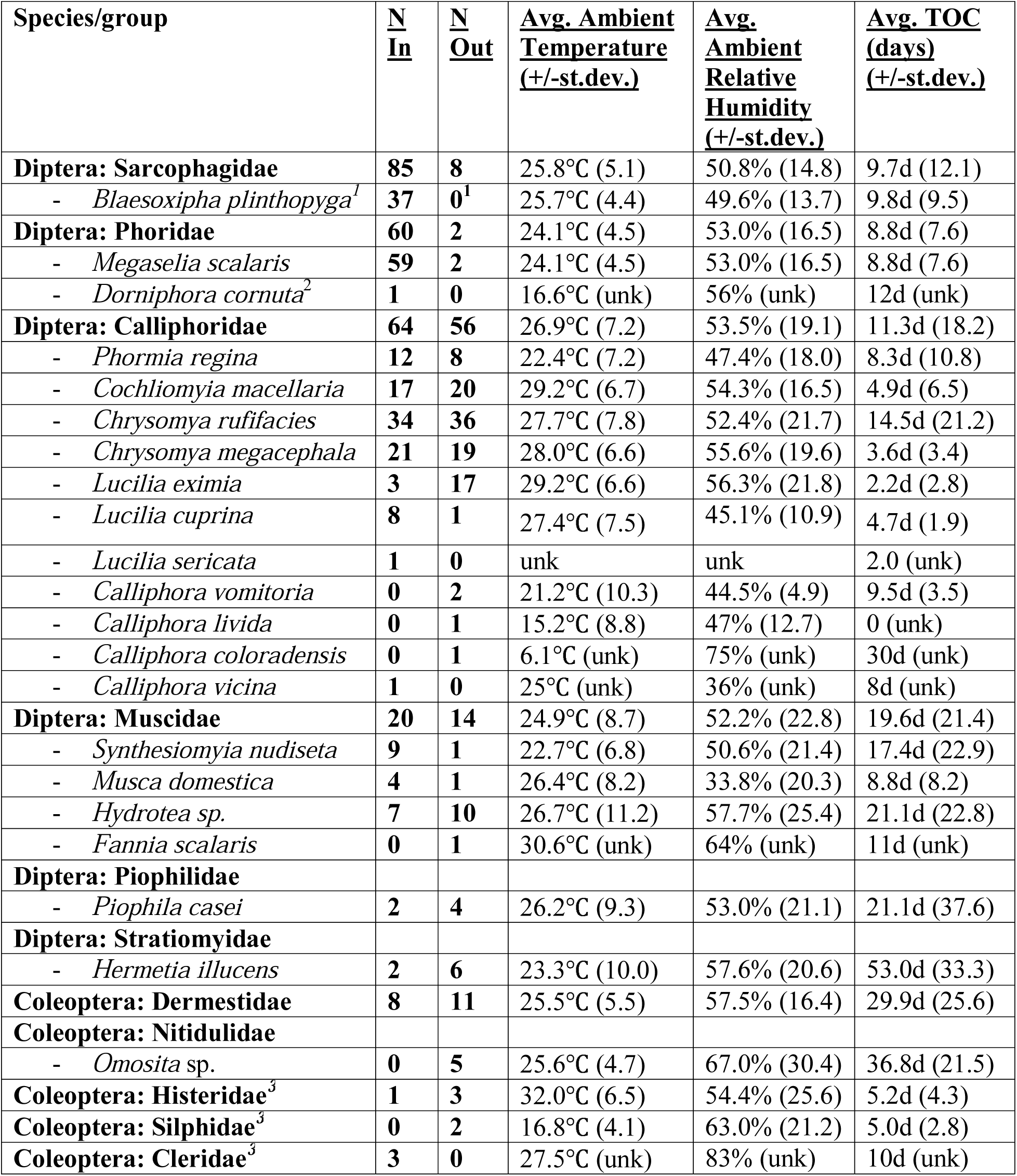
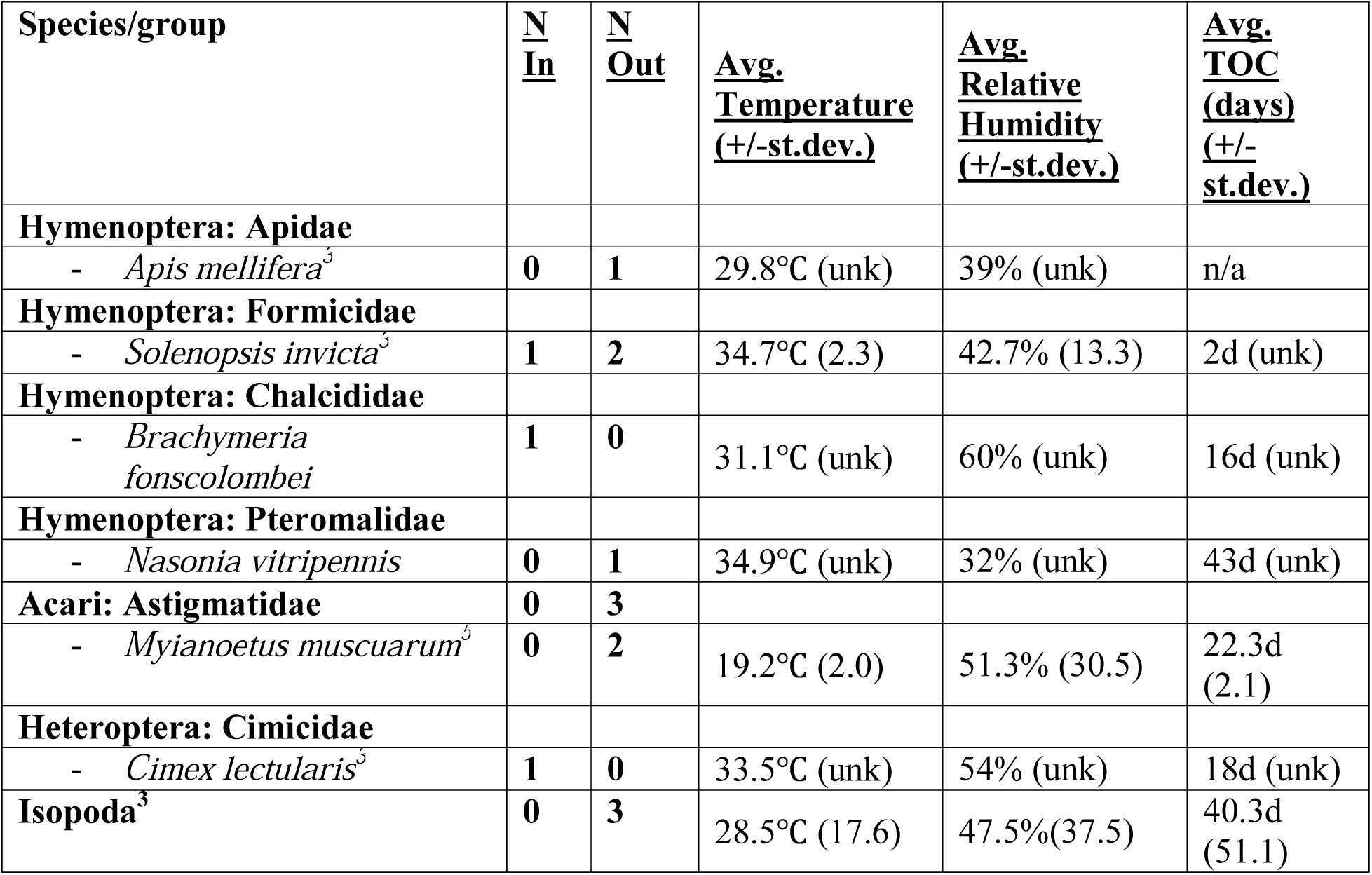
Number of cases coming from indoor and outdoor scene investigations with the average observed ambient scene temperature (℃) relative humidity (%) and the average maximum estimated time of colonization (TOC) for selected species collected in Harris County from January 2013 through April 2016 (N=203). ^1^Confirmation of this species is based on the appearance of male genitalia from reared adult males and this could be an underestimate of actual abundance for this species because reared males may not always be obtained. ^2^Identification of this specimen is based on a single larva and is tentatively identified to the species level. ^3^These species may be regularly observed at scenes but not regularly collected and reported upon because they are not typically related to time of colonization estimation. ^4^This species was recently associated with *Synthesiomyia nudiseta* [39].

### Scene Types

#### Hospitals

Insect analysis involving hospital cases typically involved the identification of insects or insect artefacts such as honeybee (*Apis mellifera* L.) stingers (Table 1) rather than TOC estimation. In those cases where a TOC estimate was generated, the cases involved perimortem myiasis wherein the decedent was colonized by insects, namely flies (Diptera; Table 1), at some time point shortly before death. Estimation of TOC for decedents that come from the hospital with fly larvae allows for the determination of colonization that can account for myiasis. Otherwise without any extenuating circumstances, such as a significant discrepancy between the stage of the insect larvae and the stage of decomposition or known medical history, myiasis can be overlooked [21]. Even when myiasis is known the calculation of the TOC estimate can be complicated by unknown factors related to which temperatures to use when calculating the accumulated degree hours (ADH) required for development [21]. This can be due to health related factors such as reduced circulation to extremities or due to unknown timing of demise and the switch from the use of body temperature to ambient temperature in calculations [24].

Appreciating the complications associated with temperature, TOC estimation using insects associated with wounds can provide useful information related to the timing of terminal events. For example if a wound, which has been colonized by insects, is associated with a fall, a TOC estimate may be a useful addition to the overall timeline of terminal events for the decedent. Timing of injuries can be difficult for individuals that have a delayed death as without scene details or witnesses the insects may provide the only timeline information available regarding injuries.

Thus far, the insects and arthropods associated with decedent’s coming from the hospital have included honeybee, typical human parasites including lice and ticks, and blow flies (Table 1). There is a growing appreciation for the information that these organisms can provide. As an example, a recent case demonstrated the potential of ticks to be useful to unraveling the travel history and potentially the next of kin (NOK) information for a decedent. In this case, a decedent came from the hospital emergency room with very little information, except that he had been traveling and had been dropped off at the hospital by a friend. The decedent had a partially engorged tick attached to his right ear that detached upon his arrival to the morgue and was quickly collected. This particular specimen was identified as a nymphal *Amblyomma* sp. most likely *A*. *cajennense* or *A*. *imitator*, which both have a known distribution in South Texas, Mexico and Central America [25]. Once all the investigative information was put together, the range for the tick and the decedent’s travel history matched. One can imagine in the absence of identification and/or NOK information that notice could be sent to consulate offices in the range of the tick to help in the search or for a decedent that is from within the United States to law enforcement agencies in the range of the tick species. One could also use the tick to test for tick-borne disease, potentially for surveillance or in determination of cause of death. Insects and other arthropods have the potential to help with a variety of other types of questions related to a decedent’s life and death; therefore, collection of specimens, no matter how trivial, can sometimes prove to be incredibly useful to the investigation.

#### Outdoor Scenes

Cases involving specimens from decedents found outdoors primarily involve estimating TOC for the purposes of estimating PMI. This is the classic scenario in forensic entomology from which many of the protocols and standards were derived [26,27]. As we integrate forensic entomology into routine medicolegal death investigations we can appreciate the breadth of opportunities available for casework and applications of insects to death investigations [6]. Figure 1 illustrates the official manner of death classifications for HCIFS forensic entomology cases from January 2013 through April 2016. It shows that outdoor scenes are outnumbered by those that occur indoors. A majority of all of the cases involve natural manner of death classifications as the official manner of death. Initially these cases come into the morgue as undetermined cases due to the difficulty of assessing trauma of decomposing decedents. The number of cases involving undetermined, accident and suicide manners of death that involve insects and occur outdoors are approximately equal to those that occur indoors. These data suggest expanding forensic entomology tools and research to understanding insect colonization indoors, which is an area of forensic entomology that has received comparatively less attention [28,29]. Noticeably, there are a higher percentage of homicide cases involving insects from decedents at scenes located outdoors than indoors. This is reflected in the initial development of forensic entomology tools for the outdoor scene as the initial opportunities of forensic entomology casework were to assist with homicide investigations [1]. It follows that decomposing decedents have fewer delays in colonization related to insect access and more opportunities for temperature modifications that may accelerate decomposition (e.g. direct sun exposure).

Analysis of factors associated with outdoor scenes in Harris County involves significantly higher average temperature (28.0°C; Table 3; Figure 2) but few other significant differences in the other factors observed when compared to indoor scenes (Table 3). The TOC estimates for outdoor scenes are slightly longer on average but not significantly different from those estimated from indoor scenes (Figure 3). Whereas the TOC estimates from hospitals were significantly shorter from both (Figure 3), which is most likely due to these cases being associated with known myiasis. Only one insect species was significantly associated with being encountered at outdoor scenes, the green bottle fly, *Lucilia eximia* (Table 4). All other insect groups, which were encountered in five or more cases, were either found to have a significantly higher probability to be encountered at indoor scenes or to be encountered at both indoor and outdoor scenes (Table 4). According to the binomial test, the probability of collecting *L*. *eximia* indoors is significantly lower than at an outdoor scene based on the data currently available in Harris County (Table 4). This suggests that finding this species on a decedent endorses outdoor colonization of the decedent when considered alone. However, three cases involved the collection of *L*. *eximia* at indoor scenes (Table 1) suggesting that under some circumstances this species does colonize bodies located indoors. The combination of species encountered on a decedent may suggest movement of the decedent if for example, when *L*. *eximia* is found together with a strongly indoor colonizing species like *M. scalaris* (Table 4), but this may only be of importance when other scene indicators suggest this possibility. The opposite scenario may also be suggested if a strongly indoor colonizing species such as *M*. *scalaris* is found on a decedent located outdoors. However, it was not analyzed with these data but the combination of species found on the decedent may temper the strong association of certain species with outdoor locations as suggested by the small number of cases where *L*. *eximia* colonized indoors. In the casework analyzed here, it was actually more rare for bodies to be found to be colonized by a single insect species with only 14.2% (28 of 197) of cases found to have only a single colonizing species and the rest with more than one.

**Table 3.** Kruskal-Wallis Chi-Squared statistical comparisons for ambient scene temperature, relative humidity and length of the TOC estimate (days) by scene location and stage of decomposition. ^1^Temperature and relative humidity comparisons did not include hospital cases, as these measurements were not made at the hospital. *indicates a significant difference at the α=0.05 level.

| Comparison | Kruskal-Wallis<br>X<sup>2</sup> Test Statistic | d.f. | P-value |
| --- | --- | --- | --- |
| Ambient scene temperature by scene location<sup>1</sup> | 8.1531 | 1 | **0.00430\*** |
| Relative humidity by scene location<sup>1</sup> | 1.0947 | 1 | 0.2954 |
| Length of TOC estimate by scene location | 7.9093 | 2 | **0.01917\*** |
| Ambient scene temperature by stage of decomposition | 2.0434 | 4 | 0.7278 |
| Scene relative humidity by stage of decomposition | 2.1554 | 4 | 0.7072 |
| Length of TOC estimate by stage of decomposition | 77.682 | 4 | **>0.0001\*** |

**Table 4.** Exact binomial test results indicating the probability of success as defined by encountering a particular insect group indoors. The 95% confidence interval and P-value for each test are also provided. *indicates a significant difference at the α=0.05 level. Statistics were only calculated for cases where each insect group was encountered five or more times during the study period.

| Groups with $\geq 5$ cases | N | Probability of success | 95% confidence interval of success | | P-value |
| --- | --- | --- | --- | --- | --- |
|  |  |  | min | max |  |
| *Blaesoxipha plinthopyga* | 39 | 0.9744 | 0.8652 | 0.9994 | **1.46E-10\*** |
| *Megaselia scalaris* | 67 | 0.9701 | 0.8963 | 0.9964 | **<2.20E-16\*** |
| *Lucilia cuprina* | 11 | 0.9091 | 0.5872 | 0.9977 | **0.01172\*** |
| Sarcophagidae | 103 | 0.9029 | 0.8287 | 0.9525 | **<2.20E-16\*** |
| *Synthesiomyia nudiseta* | 18 | 0.8889 | 0.6529 | 0.9862 | **0.001312\*** |
| *Musca domestica* | 5 | 0.8000 | 0.2836 | 0.9949 | 0.375 |
| *Phormia regina* | 31 | 0.5806 | 0.3908 | 0.7545 | 0.4731 |
| Calliphoridae | 134 | 0.5448 | 0.4565 | 0.6310 | 0.342 |
| *Chrysomya rufifacies* | 78 | 0.5000 | 0.3846 | 0.6154 | 1 |
| *Cochliomyia macellaria* | 46 | 0.4783 | 0.3289 | 0.6305 | 0.883 |
| *Hydrotea* sp. | 21 | 0.4762 | 0.2571 | 0.7022 | 1 |
| *Dermestes maculatus* | 22 | 0.4091 | 0.2071 | 0.6365 | 0.5235 |
| *Piophilidae casei* | 7 | 0.2857 | 0.0367 | 0.7096 | 0.4531 |
| *Hermetia illucens* | 8 | 0.2500 | 0.0319 | 0.6509 | 0.2891 |
| *Lucilia eximia* | 22 | 0.0000 | 0.0000 | 0.1544 | **4.77E-07\*** |
| Nitidulidae | 5 | 0.0000 | 0.0000 | 0.5218 | 0.0625 |

**Figure 2:**
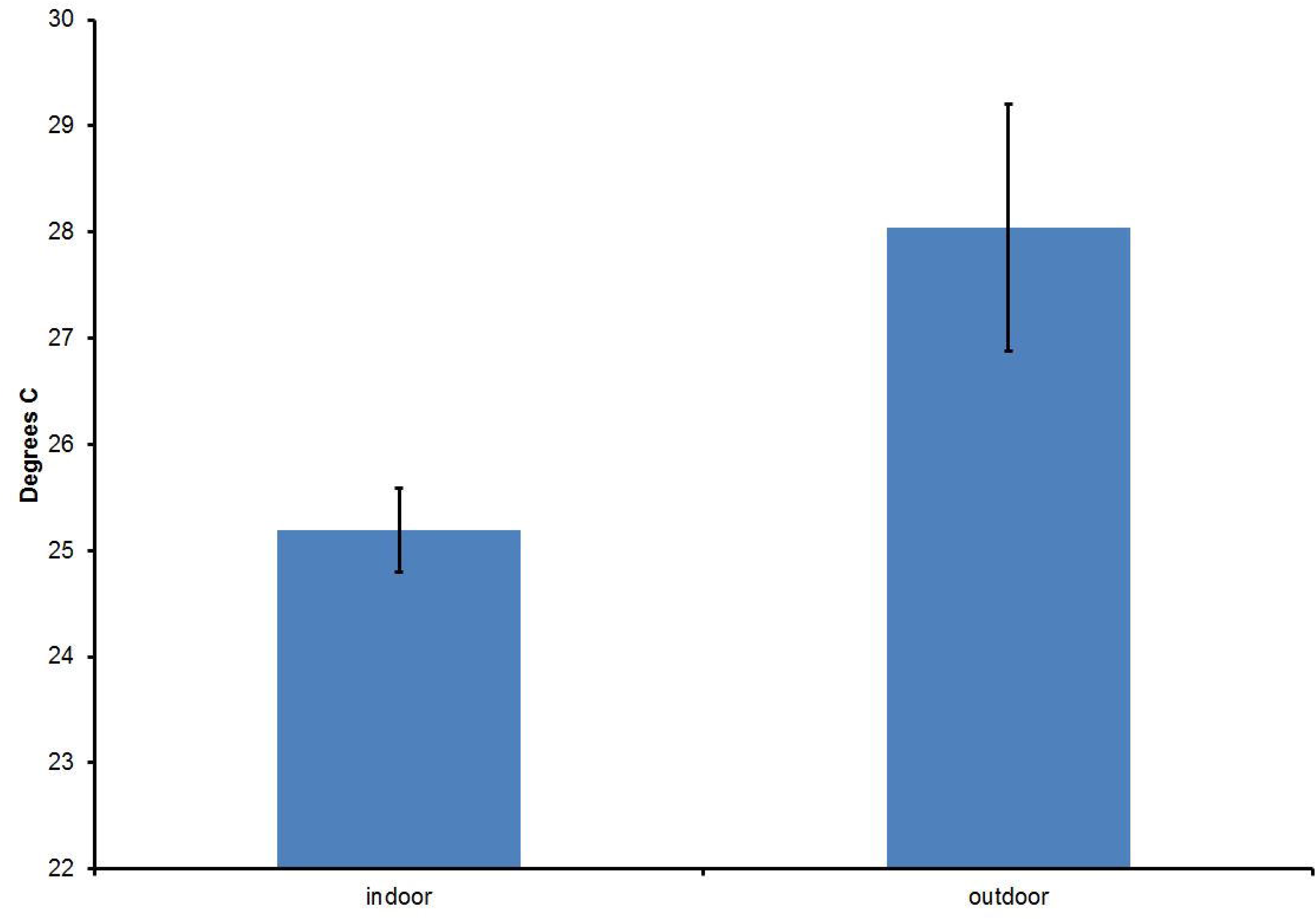
Average ambient scene temperature by scene location. Average (+/-standard deviation) ambient air temperature (°C) recorded near the decedent at scenes located indoors and outdoors for forensic entomology cases over the study period (January 2013 – April 2016). (N=203).

**Figure 3:**
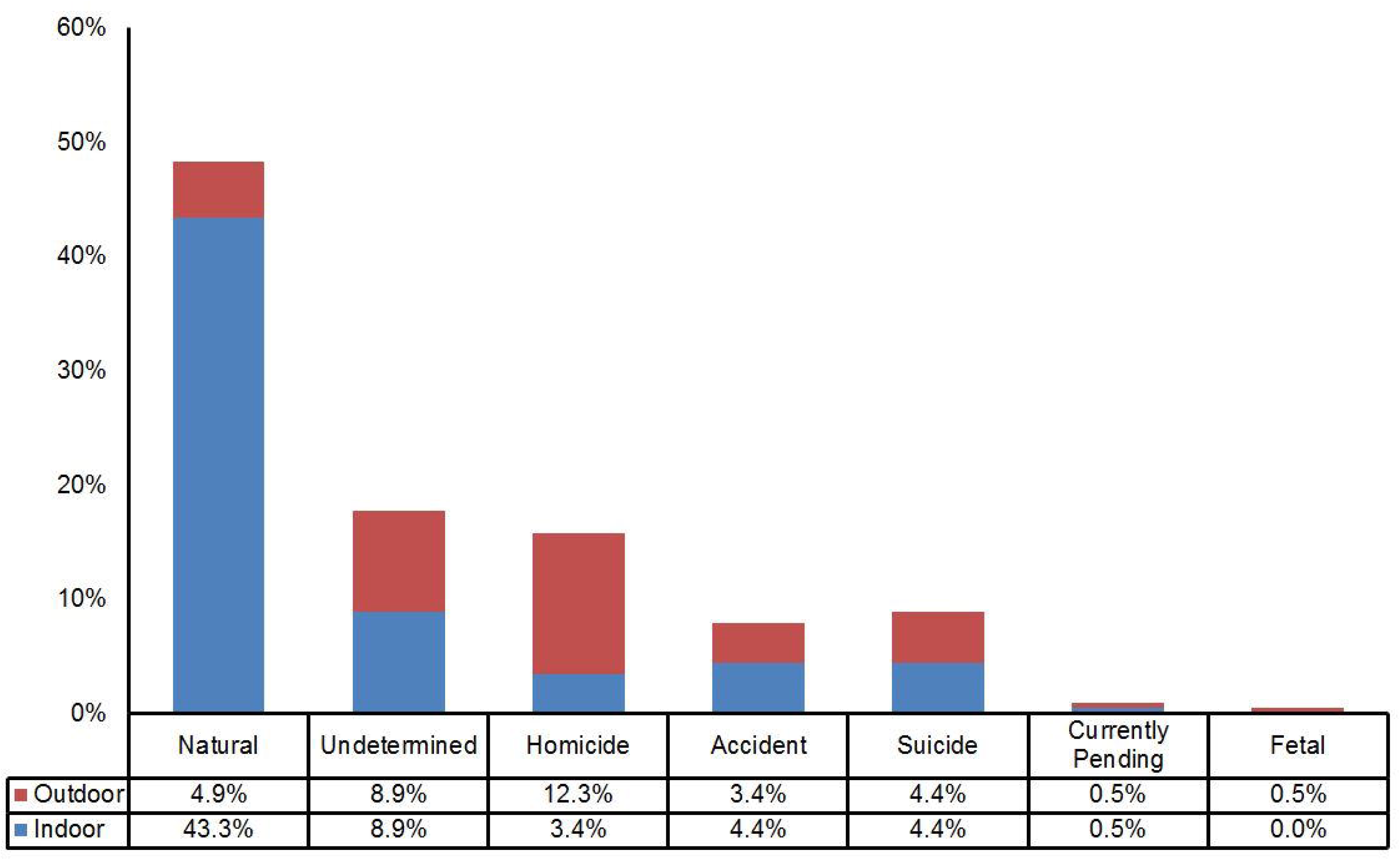
Average TOC length (days) by scene location. Average (+/-standard deviation) time of colonization (TOC) estimate (days) by scene location for forensic entomology cases over the study period (January 2013 – April 2016). (N=203).

#### Indoor Scenes

Cases involving insects collected indoors also have a primary goal of TOC estimation for the purpose of estimating PMI. A majority of the forensic entomology case reports from Harris County over the analyzed period were from indoor scenes and they encompassed every manner of death classification except fetal (Figure 1). The average ambient scene temperature observed at indoor scenes (25.2°C) was significantly lower than that observed at outdoor scenes (Table 3; Figure 2). As mentioned earlier the number of cases that involve manner of death classifications involving accidents, suicides and undetermined classifications involve scenes located indoors and outdoors suggesting a need for more tools for indoor scenes. Indoor scenes can be complicated by multiple factors that affect insect colonization (e.g. delays in colonization; [28]), temperature modification (e.g. air conditioning, heating), co-located decomposing remains (e.g. decomposing pets; [30]), 24-hour lighting which may allow for nocturnal oviposition (e.g. a TV or lamp left on at the scene;[31]), other sources of insects in the residence that may reduce or eliminate a delay in colonization (e.g. hoarding of trash and food) and a number of other case specific factors that may affect the TOC estimate. Many of these complicating factors have as yet unknown effects on TOC estimation but have the potential to have significant impacts on PMI estimation.

Another significant observation regarding indoor scenes is in the different insects found at indoor scenes. Just as some insect species appear to have preference for outdoor scenes there appears to be a strong association between indoor scenes and several insect groups including *Megaselia scalaris*, Sarcophagidae including *Blaesoxipha plinthopyga*, *Synthesiomyia nudiseta* and *Lucilia cuprina* as can be observed in the binomial test statistics presented in Table 4. This suggests when each of these species, considered alone, may represent an indicator of indoor colonization or at the very least allow for the consideration of an indoor scene from the insect’s perspective. A car in a covered carport or a storage container where a decedent was living may not be what may typically be considered indoor scenes but from the insect’s perspective, they may fulfill colonization preferences. Therefore, scene indicators and context should all be taken into account when considering insect specimens as indicators of body movement.

These results are similar to those observed by other researchers in other geographic locations. On the Hawaiian Island of Oahu, Goff [32] found that three species were strong indicators of indoor scenes including a species of the Phoridae *Megaselia scalaris*, the Sarcophagidae *Bercaea haemorrhoidalis* and the Muscidae *Stomoxys calcitrans*. While not exactly the same species encountered in the current study, a similar combination of representatives of the same fly families was found with the same Phoridae *M*. *scalaris*, the Sarcophagidae *B. plinthopyga* and the Muscidae *S*. *nudiseta* indicating indoor colonization. Goff also found that there was a wider diversity of fly species indoors at least preliminarily. In the cases analyzed in this study there was a similar, though small, result with an average of 1.9 (+/- 1.3 standard deviation) species of flies found from outdoor scenes and an average of 2.4 (+/-1.3 standard deviation) fly species encountered in specimens from outdoor scenes overall. The Hawaiian study also reported a higher diversity of beetles encountered outdoors and this was also a trend that was observed in the current study although beetles were more rarely collected (Table 1). In Western Canada Anderson [28] found that most of the blow fly species colonizing experimentally placed pigs indoors and outdoors overlapped with several species appearing to be restricted to the outdoors. Similar results were obtained in this study where the only species that was strongly associated with outdoor scenes was *L*. *eximia* (Table 1 and 4). However, in an analysis of casework Anderson [18] found that in British Colombia, Sarcophagidae and Phoridae did not appear to be significant colonizers and did not appear to have the same dichotomy based on scene location. When one considers the climate similarities and differences among the three geographic locations, such as the temperatures, relative humidity (Table 2) and precipitation encountered at scenes in Harris County along the Gulf Coast are perhaps more similar to those encountered in Hawaii rather than British Columbia. Furthermore climate change may lead to changes in species ranges over time [33,34] or to differentiation in developmental progression in different populations of the same species. This highlights the necessity and importance of collecting and publishing these data for different geographic locations, different climatic regions and different species assemblages rather than relying only on a handful of studies.

Some species of insects and arthropods analyzed in Harris County casework have only been found indoors thus far. The flesh fly *Blaesoxipha plinthopyga* is the only fly, which has been identified exclusively indoors. This fly was not known to be a forensically significant fly in the US until quite recently [35] and this is perhaps due in part to the difficulty in identification of Sarcophagidae, which requires either prepared adult male flies [36] and expert morphological expertise associated with male genitalia or expertise in the molecular systematics of Sarcophagidae [37,38] and adequate laboratory capacity. Therefore, Sarcophagidae have often been relegated to the Family level without further identification. Nevertheless, even with identification there are significant gaps in the literature for detailed development data sets for many of the forensically important Sarcophagidae species. The mite, *Myianoetus muscuarum*, has only been found indoors but it has only been found in a few cases thus far ([39] and Table 1) and it may be the case that the more it is looked for the more it may be found. The other species that have been found exclusively indoors have very low case numbers and may have just not been collected frequently enough to establish habitat or colonization preferences for these insects/arthropods (Table 1).

### Insects/Arthropods

The introduction of routine forensic entomology into the Medical Examiner’s office has allowed for the observation of several new associations between insects, arthropods and human decomposition. One of those observations was the first report of what was determined to be myiasis of a decedent shortly before his death by *L*. *eximia* and *C*. *rufifacies* for the first time on a human in the US [40]. In another case, the blue blow fly, *Calliphora coloradensis* was collected from an outdoor scene in February 2014, associated with a decedent in an advanced state of decomposition and vertebrate scavenging. The specimen was reared from a pupa located next to the decedent and it was the first time that this species had been collected from a decedent. It is known to occur in Texas [15] but has not to the author’s knowledge, been collected in association with a decomposing human in Harris County. It has been previously recorded as an adult associated with a human corpse found at high elevation in Colorado but was not observed as a colonizer of the remains [41]. In another case the tawny crazy ant, *Nylanderia fulva* was recorded on a decomposing human body in Harris County for what appears to be the first time (Table 1).

Aside from new insect associations, other new arthropod associations are being made as well. The mite, *M*. *muscuarum* was recently established to have a relationship with *S*. *nudiseta* at indoor scenes [39] and has been found in additional cases since this publication was completed. Mites continue to be encountered at scenes and with help of Dr. B. OConnor (University of Michigan), identifications of the mite species are being made in order to develop a taxonomic index and to be able to associate the mite fauna with timing in the decomposition process for this geographic location. A nematode was recently found in association with decomposing decedents found in advanced stages of decomposition located indoors. These nematodes were first discovered in a rearing container from a case involving a decedent found in an advanced state of decomposition in a hoarder home. The decedent was also colonized by Sarcophagidae, Calliphoridae and *Hermetia illucens*. Rearing of the specimens to confirm identifications led to the discovery of large numbers of nematodes in the rearing container. The identity of these nematodes was difficult as they may represent a new species and their identity is still being investigated by Dr. Y. Qing (Canadian National Collection of Nematodes).

The location of Harris county being on the Gulf Coast and near the southern US border presents the opportunity to make observations related to new species introductions as well as documenting new associations. The forensic entomologist can therefore serve in a capacity to identify foreign species introductions and to notify relevant interested parties. This applies to not only those insects that are used for TOC estimation but also those that may be relevant to public health and cause of death determination as new arthropod-borne diseases are introduced to the US.

### Seasonality

The seasonality of the blow flies encountered in Harris County follows patterns generally accepted for Calliphoridae in North America [42]. The typical cool weather blue bottle flies, *Calliphora* spp. are found in the winter months and other species such as the black blow fly, *P*. *regina*, and the Muscidae species, *S*. *nudiseta* are active in the spring months (Table 1). The green/bronze bottle flies, *Lucilia* spp. are most often collected in the hot summer months but can be encountered throughout the spring through the fall months (Table 1). The introduced species, the hairy maggot blow fly, *Chrysomya rufifacies* and the oriental latrine fly, *Chrysomya megacephala* appear to be active year-round in some years whereas in other years they can be absent in the winter months (Table 1). It is a very rare occurrence for there to be snow in Harris County and temperatures below 0.0℃ rarely persist for more than a few days if at all [43]. These climate characteristics probably help the introduced tropical Calliphoridae species persist through the winter as freezing temperatures are thought to restrict their distribution in North America [44].

The Sarcophagidae species, *B*. *plinthopyga*, and the Phoridae species, *M*. *scalaris*, are not as seasonally restricted and appear to be collected in casework year-round (Table 1). It would seem that this might be partially explained by their exploitation of indoor scenes. Currently no data is available on the dispersal patterns of either of these fly species. Other Phoridae species appear to be capable of transportation via wind and human modes of transport [45]. Preliminary data suggests population structure exists within *M*. *scalaris* populations in the US even across relatively small geographic distances [46]. Indoor environments with their cooler and less variable temperature range (Figure 2) may offer refuge when outdoor conditions are unfavorable allowing populations to persist locally during bad weather.

## CONCLUSIONS

The integration of full time forensic entomology services to the medical examiner’s office in Harris County has revealed new observations, confirmed long-standing hypotheses and opened up a wide range of new opportunities. The casework described here is not an endpoint and new species observations and associations continue to be revealed. This study represents a reference point from which to describe additional observations and trends and from which to compare other geographic locations and species assemblages. These data also remind us that depending upon generalizations and assumptions can be misleading and that keeping an open mind and collecting something that may at first seem trivial may prove to be exceptionally useful.

The data presented here also have illustrated several persistent issues in the application of forensic entomology to casework. While molecular identification tools are being more widely developed and applied to morphologically difficult groups (e.g. Sarcophagidae and Muscidae) there are significant gaps in development data sets. A single development data set is available for *B*. *plinthopyga* which was never intended for use in application to forensic entomology as it is includes total generation time for specimens collected during a study of diapause [47]. Yet this species is considerably important to casework in Harris County (Table 1). There are no known published development data sets for *C*. *coloradensis*, *C*. *livida* or *D*. *maculatus* using a diet that approximates human tissue. The suggested practice of using closely related species to approximate development for those species that have missing data introduces unnecessary uncertainty in an estimate of TOC and its application to PMI. Furthermore there is growing awareness that different populations of the same species have different developmental responses to the same temperatures over large [48] and relatively small geographic distances [49]. This strongly suggests that using development datasets from populations not local to the collection location for the case may introduce unwelcome uncertainty to a case that becomes difficult to track and account for once the case has been filed. Several of the species that are common and important to casework in Harris County lack local development data sets including *C*. *megacephala*, *S*. *nudiseta*, *L*. *eximia* and *L*. *cuprina*.

While missing development datasets can be generated, other issues continue to haunt forensic entomology casework that are not and will not be as easy to remedy. In 1992, Catts [50] outlined several complications in using insects to estimate postmortem interval. These issues, such as maggot mass heat, insect access and entomotoxicology, continue to be sources of uncertainty for the application of forensic entomology to casework. It has only been relatively recently that there has been an increased awareness for the need to account for uncertainty and sources of error in forensic entomology within the context of all forensic sciences in the US [51]. In addition, while identifying potential sources of error is more widely appreciated accounting for how these errors and influences impact individual TOC and PMI estimates has not yet been resolved. Tackling these challenges in forensic entomology will require strong collaboration between academic and practitioner parts of the community but the results will make forensic entomology an even more powerful tool in understanding the post-mortem interval.

## ACKNOWLEDGEMENTS

The author would like to acknowledge the help of Drs. Barry OConnor (Acari; University of Michigan), Terry Whitworth (Calliphoridae; University of Washington), Martin B. Berg (Chironomidae; Loyola University), Robert Puckett (Formicidae; Texas A&M University), Pete Teel (Ixodidae; Texas A&M University), Qing Yu (Nematoda; Canadian National Collection of Nematodes), Brian Brown (Phoridae; Natural History Museum of Los Angeles) and Greg Dahlem (Sarcophagidae; Northern Kentucky University) for their generous assistance with identification of specimens from their respective taxonomic specialties. The help of these taxonomic experts adds valuable taxonomic information to not only casework reports but to the body of knowledge regarding insects and associated arthropods found on decomposing human remains. The author also wishes to thank Drs. Jeffery Tomberlin, Adrienne Brundage, Micah Flores and Gail Anderson for their assistance in implementing and maintaining 100% outside peer review of forensic entomology casework for the Harris County Institute of Forensic Sciences (HCIFS). This could not have been accomplished without their gracious voluntary service to this effort. The author also wishes to acknowledge the forensic pathologists and forensic investigators of the HCIFS for their support in embracing insects in their day-to-day activities. The integration of forensic entomology into the daily activities of the medical examiner’s office has greatly benefitted the field and will continue to do so in the future.

